# Robust genome editing with short single-stranded and long, partially single-stranded DNA donors in *C. elegans*

**DOI:** 10.1101/352260

**Authors:** Gregoriy A. Dokshin, Krishna S. Ghanta, Katherine M. Piscopo, Craig C. Mello

## Abstract

CRISPR-based genome editing using ribonucleoprotein (RNP) complexes and synthetic single stranded oligodeoxynucleotide (ssODN) donors can be highly effective. However, reproducibility can vary, and precise, targeted integration of longer constructs – such as green fluorescent protein (GFP) tags remains challenging in many systems. Here we describe a streamlined and optimized editing protocol for the nematode *C. elegans*. We demonstrate its efficacy, flexibility, and cost-effectiveness by affinity-tagging all twelve of the Worm-specific Argonaute (WAGO) proteins in *C. elegans* using ssODN donors. In addition, we describe a novel PCR-based partially single-stranded “hybrid” donor design that yields high efficiency editing with large (kilobase-scale) constructs. We use these hybrid donors to introduce fluorescent protein tags into multiple loci achieving editing efficiencies that approach those previously obtained only with much shorter ssODN donors. The principals and strategies described here are likely to translate to other systems and should allow researchers to reproducibly and efficiently obtain both long and short precision genome edits.

## Introduction

In theory, CRISPR/Cas9-based genome editing enables researchers to rapidly generate designer alleles of any locus for genetic, cytological, or biochemical analyses. In practice, however, we have found that the technology remains far from routine for many users, especially in applications where long templated insertions are desired. Here we describe a streamlined and optimized protocol for genome editing the nematode *C. elegans*. We find that seemingly minor details, such as the order of donor DNA addition to the editing mix, can have dramatic consequences for editing efficiency. We demonstrate pronounced toxicity of RNP at high concentrations and provide a strategy for optimizing RNP levels using a co-injected, easily scored reporter. Finally, we show that generating hybrid, partially single stranded long DNA donor molecules dramatically promotes templated repair for the insertion of longer edits such as GFP. Although, we have only tested these strategies in *C. elegans*, it seems likely that the principals revealed here including the following key features, will be relevant in other systems:

- Utilization of a robust, Cas9-independent co-injection marker to control for injection quality, to optimize Cas9 RNP concentration, and to monitor toxicity.
- Pre-assembly of the RNP complexes before adding donor DNA to the injection mixture.
- Employment of hybrid PCR-based donors with single-stranded homology arms for consistent, high-efficiency insertion of large constructs.

## Results

### Cas9 RNPs are toxic at high concentrations

In the course of adopting Cas9 RNP editing methodologies (PAIX *et al.* 2015) we decided to monitor injection quality by adding the *rol-6(su1006)* plasmid to the injection cocktail (MELLO *et al.* 1991). We were very surprised to find that, despite giving high numbers of edited progeny, the numbers of transgenic Roller (*rol-6*) animals were greatly reduced. We noted that the few surviving Roller animals obtained were often sick and sterile, suggesting that toxicity, or excessive genome editing might cause the lack of Roller transgenics (data not shown).

To address these possibilities, we performed a titration of RNP concentrations while holding the Roller DNA concentration constant. We then examined both the genome editing efficiency and the frequency of Roller transgenics among F1 progeny of the injected animals. Worms expressing the bright fluorescence marker GFP::GLH-1 were co-injected with 40ng/μl pRF4::rol-6(su1006) plasmid and dilutions of Cas9 RNPs loaded with an anti-gfp guide (Figure 1A). In our pilot studies we recovered very few Rollers at 2.5μg/μl of Cas9 used in initial *C. elegans* Cas9 RNP protocols (CHO *et al.* 2013; PAIX *et al.* 2015), we therefore decided to begin with a 5 fold dilution, 0.5 μg/μl as a starting RNP concentration. This concentration was recently proposed by Prior and colleagues (PRIOR *et al.* 2017). Injections using 0.5 μg/μl of Cas9 resulted in an average of 17 F1 roller progeny per injected P0 animal. Reducing the concentration by 2-fold, down to 0.25μg/μl doubled the frequency of F1 rollers to 33, while a ten-fold dilution to 0.025μg/μl resulted in 43 F1 roller progeny per P0 (Figure 1B). These latter two F1 roller frequencies are comparable to rates reported for pRF4::rol-6(su1006) injected alone (MELLO *et al.* 1991). Taken together these findings suggest that RNP concentrations below 0.25 μg/μl do not interfere with expression of the co-injected Roller marker gene.

**Figure 1.**
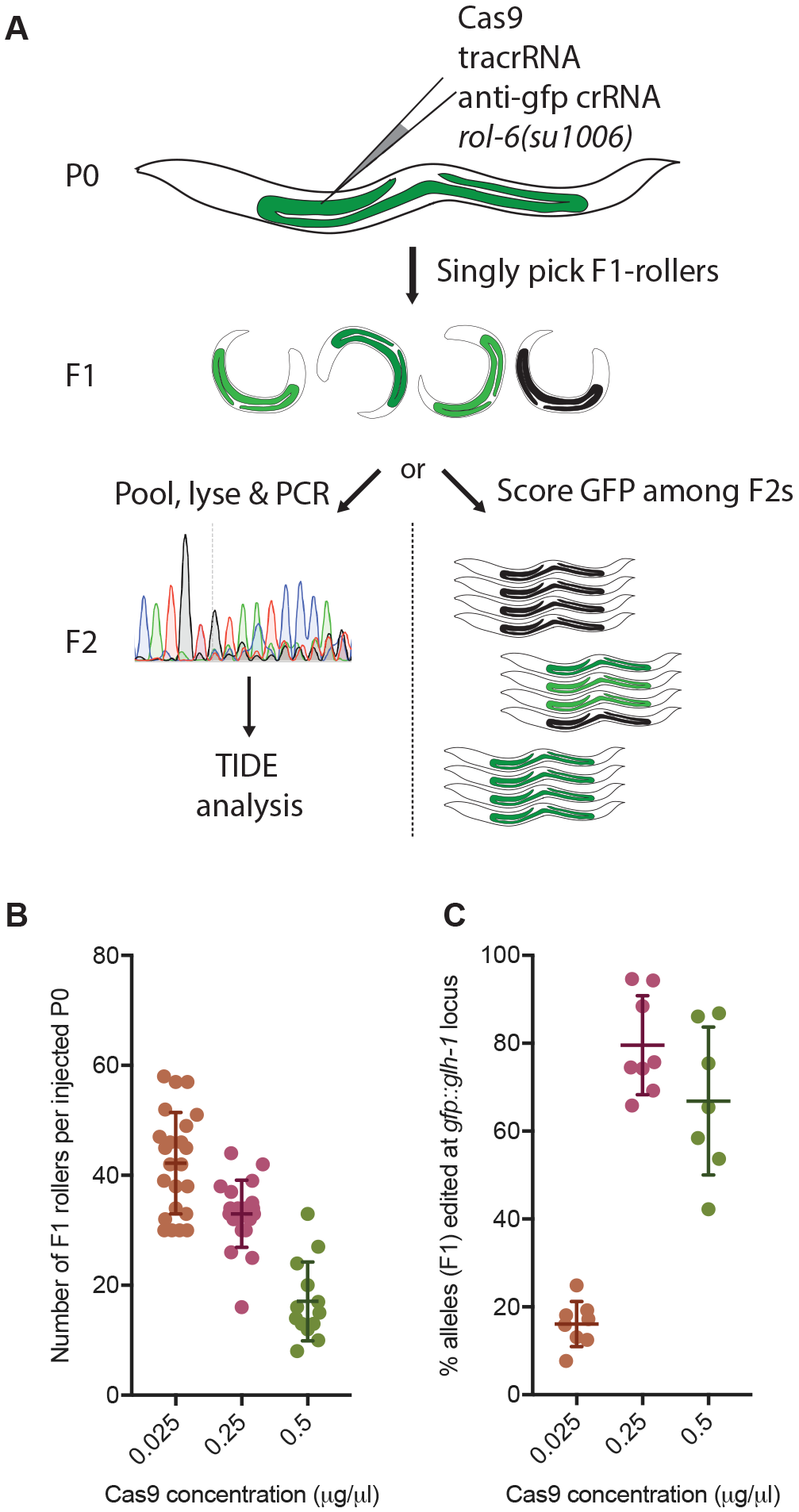
Determining optimal Cas9 RNP concentrations. (**A**) Schematic representation of the optimization workflow. Cas9 protein loaded with anti-GFP guide is co-injected at several concentrations with 40ng/μl of *rol-6* plasmid into *gfp::glh-1* animals. Number of F1 Rollers segregated by each injected P0 is scored. F1 Rollers are then subject to genotyping as a pool by TIDE analysis (left) or their F2 progeny are scored by microscopy (right). (**B**) Number of F1 Rollers recovered from an injected P0 at three different Cas9 concentrations. Each dot represents an individual animal. (**C**) Percent of alleles carrying an in-del at the *gfp::glh-1* locus at three different Cas9 concentrations as determined by TIDE analysis. Each dot represents a pool of F1 Rollers from one injected P0.

We next asked how Cas9 RNP concentrations affected the in-del frequency at the *gfp::glh-1* locus. To measure in-del rates in a high throughput fashion we used the TIDE analysis pipeline, which estimates the in-del rates in a mixture of PCR products (Figure 1A, left) (BRINKMAN *et al.* 2014). To do this we PCR amplified the *gfp::glh-1* locus from pools of all F1 rollers segregated by a given injected P0 worm and subjected the mixture to sanger sequencing and TIDE analysis. Using this approach we found that at 0.025 μg/μl ~16% of alleles carried an in-del. The number of edited alleles increased to ~80% at 0.25μg/μl (Figure 1C), but did not increase further when the Cas9 concentration was doubled to 0.5μg/μl, and in fact appeared to decline slightly to ~67% (Figure 1C). Because GFP::GLH-1 is easily detectable in adult animals under the florescent dissection microscope, we were able to validate the TIDE results directly using microscopy (Figure 1A, right). For example, we determined that at 0.25μg/μl of Cas9 ~98% of all F1 rollers segregated GFP-negative (successfully edited) progeny (Figure S1A). Furthermore, ~68% were homozygous, producing only GFP negative progeny, while another ~30% were heterozygous (Figure S1B). Based on these numbers we can calculate that 83% of all *gfp::glh-1* alleles were successfully edited at 0.25μg/μl of Cas9 (Figure S1C). These numbers correlate well with TIDE data (Figure 1B), and thus lend confidence to the calculations of the percentage of *gfp::glh-1* alleles cleaved at each Cas9 concentration (Figures 1C and S1C). Finally, to determine the reproducibility of these findings we repeated the injections with a previously characterized moderate-efficiency guide targeting the *unc-22* locus (KIM *et al.* 2014) and observed similar results (data not shown).

### Efficient editing with ssODN donors using a Roller plasmid co-injection marker

The above findings demonstrate that Roller plasmid co-injection identifies animals that are highly likely to undergo CRISPR-induced DNA double strand breaks. We next wished to test this methodology for achieving homology-directed repair (HDR). To do this we decided to introduce a (3X)FLAG-affinity tag into each of the twelve worm-specific Argonautes (WAGOs). For each gene we designed guides targeting the PAM site closest to the ATG start codon (without any further optimization or guide testing) (Figure 2A; Appendix 1 for guide and ssODN donor sequences) (PAIX *et al.* 2015). *wago-1* and *wago-2* are highly similar near the ATG and no specific guide could be designed; thus, one guide targeting both loci was used.) Injection mixtures were prepared by simultaneously mixing Cas9 with the guide RNA, plasmid and donor ssODN, without pre-incubating to assemble the RNP (Figure 2B). (Note: we now, instead, recommend pre-assembling the RNP prior to adding the donor DNA and plasmid (see below), as this simple change in the order of addition dramatically improved repair with longer templates. Each mixture was then injected into ~10 P0 N2 worms using standard worm gonadal injection methodology (MELLO AND FIRE 1995). Individual F1 Rollers were singled to plates and after producing broods were genotyped for 3XFLAG insertions (Figure 2B, See Appendix 2 for detailed protocol).

**Figure 2.**
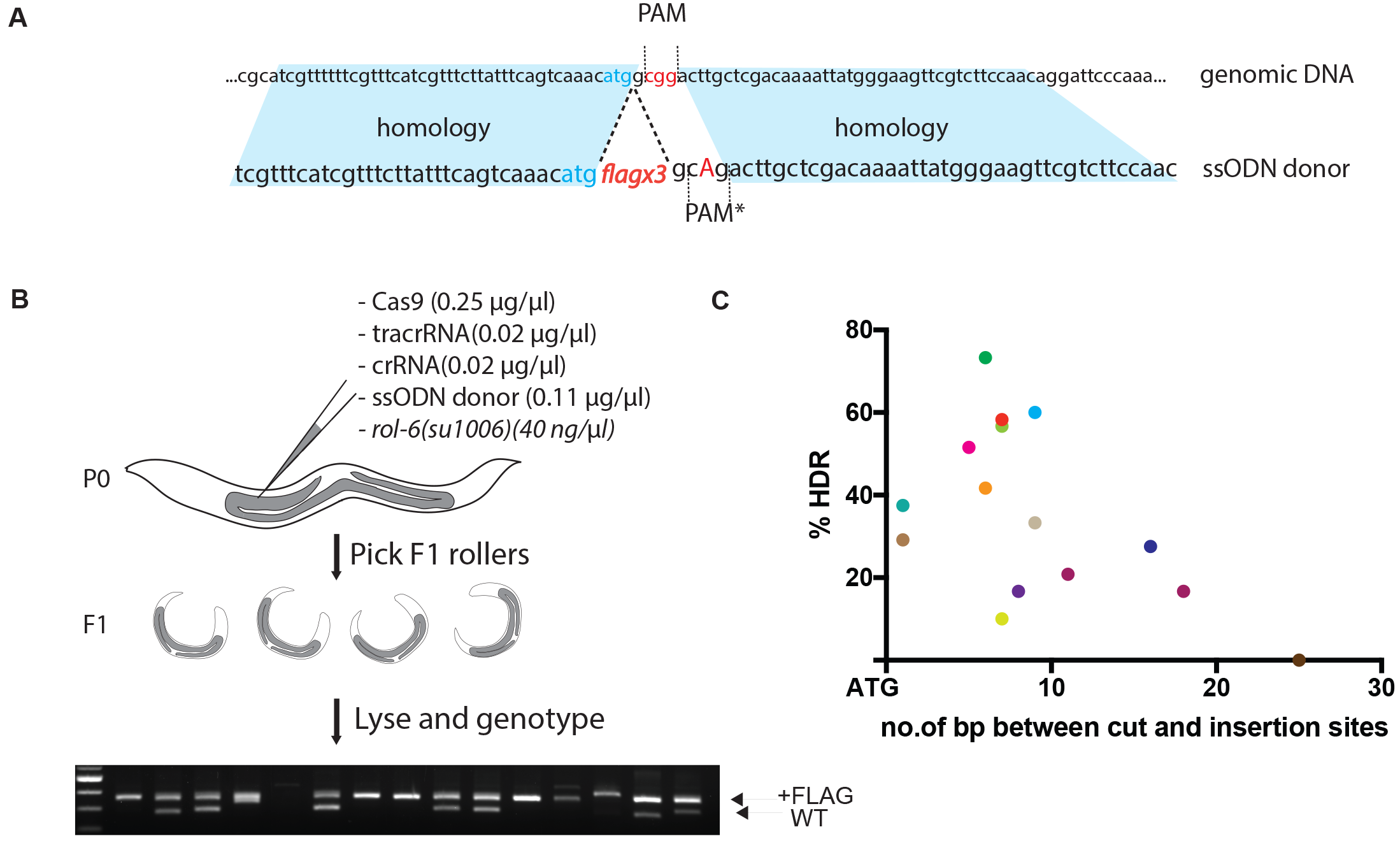
Efficient integration of 3XFLAG donor at all twelve WAGO genes with *rol-6* coinjection marker. (**A**) Schematic of donor design for 3xflag insertion directly downstream of the ATG (based on (PAIX *et al.* 2015). Blue shading highlights homology arms, red letters indicate the PAM site, blue letters represent the START codon, capital A is the mutation introduced to disrupt the PAM site in the donor. (**B**) Schematic of the CRISPR protocol. Simplified injection mixture contains just the RNP components, the ssODN donor, and *rol-6* plasmid. Approximately 24 F1 rollers from two best injection plates were single cloned and genotyped. Lower band is the wild-type PCR product; upper band is upshifted due to 3×flag insertion. (**C**) Efficiencies of 3×flag insertion plotted versus distance of the guide cut site from the START codon. Each dot represents targeting of one WAGO gene. Error bars represent standard deviation from the mean.

We were able to recover 11 out of 12 WAGO tagged strains among the first 24 F1 progeny screened by PCR from each set of injections, and in every case we recovered multiple independent alleles. The average success rate for precise insertion of the 3XFLAG tag ranged from 10-73% and averaged ~34%. Plotting the insertion efficiency versus the distance between the Cas9-induced cut and the desired insertion site (directly after the ATG start codon [Figure 2A]) we found no strong correlation up to 20 base pairs away (Figure 2C). The *wago-6* (*sago-2*) locus was the only outlier, likely because the nearest available cut site for use with the original donor design was 27 bases away from the site of insertion. Although a number of insertions were recovered at this locus they were either out of frame or contained random DNA sequences (data not shown). The *wago-6* gene contains a second PAM site located right at the ATG start codon. This site was not used originally because the 3XFLAG donor sequence (which starts with a “G”) would not disrupt the PAM. Moreover, the alternative approach to prevent re-cutting of the repaired locus, mutating the guide binding site, would require introducing potentially undesirable mutations into the 5’ UTR. To solve this problem, we added an extra CCC, proline codon, to the 3×FLAG donor sequence, immediately downstream of ATG (Figure S2). Using this donor and guide we recovered *flag::wago-6* alleles in 52% of the F1 Roller animals analyzed. In all of the edited strains the Roller phenotype was expressed only transiently during the F1. These findings demonstrate the general utility of the Roller marker for identifying edited animals without introducing additional edits or undesired phenotypes into the resulting strains. In addition, these findings indicate that as long as the desired insertion site resides within 20 bp of the cut site, ssODN donors provide for highly efficient editing.

### Hybrid dsDNA donors promote the integration of large constructs

High rates of HDR have been reported using PCR-generated double stranded DNA (dsDNA) ~1kb-sized donors with ~35bp homology arms (PAIX *et al.* 2015). However, we have struggled to reproduce these successes using the original or optimized protocols (data not shown). Extending the homology arms from 35bp to 120bp resulted in low, ~1%, but reproducible levels of GFP integration at 3 different loci (Table 1). We speculated that perhaps the additional donor dsDNA (which was 5× greater in concentration than the Roller plasmid) was interfering with Cas9 RNP assembly. We therefore decided to pre-assemble the Cas9 RNPs for 10 minutes at 37°C *prior* to adding the donor DNA or *rol-6* marker plasmid. This change in the order of addition of editing components gave a consistent increase in editing efficiency to ~4% (see Figure 3C). Nevertheless, there was still a very large gap between the efficiency of templated repair using ssODNs and longer dsDNA donors.

**Table 1:** HDR efficiencies of GFP insertion with blunt-ended PCRs as donors. All the donors consist of 120bp long homology arms on both ends.

| Locus | no. of F1 rollers (*rol-6*) |  |
| --- | --- | --- |
|  | Screened | GFP+ |
| *wago-4* | 54 | 1 (1.85%) |
| *sago-2* | 119 | 1 (0.84%) |
| *ppw-1* | 76 | 1 (1.32%) |

**Figure 3.**
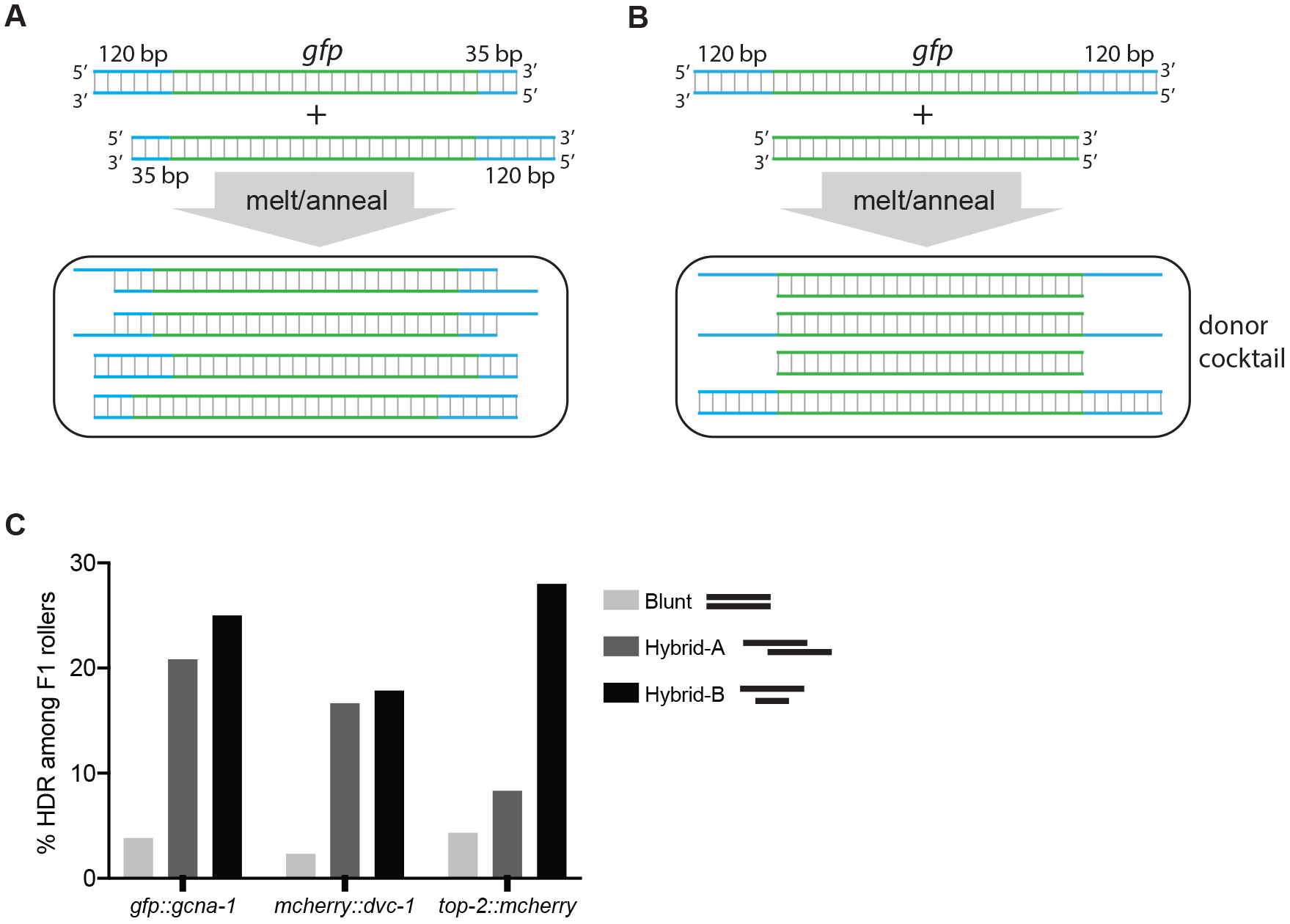
Efficient editing with long, partially single-stranded dsDNA donors. (**A**) and (**B**) Schematics of the strategy for generating hybrid dsDNA donor cocktail featuring molecules with ssDNA overhangs. (**C**) Integration efficiencies of GFP or mCherry fluorescent tags using blunt donors or hybrid dsDNA donor cocktail at diverse loci, plotted as a fraction of F1 rollers positive for appropriate insert as detected by PCR.

A recent study proposed that ssODN donors are integrated by a highly efficient single stranded template repair (SSTR) pathway, while dsDNA donors rely on a less efficient homology-directed repair (HDR) pathways (RICHARDSON *et al.* 2017). We therefore wondered whether we could achieve the improved efficiency of ssDNA by employing large PCR-based donors with single stranded overhangs. To test this idea, we generated two PCR donors to target the same locus: one with a 120bp left homology arm and a 35bp right homology arm, and the other with 35bp on the left and 120bp on the right. By mixing these donors at equimolar quantities, then melting and re-annealing the mixture we should get a mixture of four different molecules (Figure 3A) two of which have either 3′ or 5′ single stranded overhangs. Alternatively, hybrid asymmetric PCR donors were prepared by annealing molecules with 120bp homology arms to a PCR product containing just the insert, with no homology arms (Figure 3B). 200ng/μl of blunt donor or hybrid cocktail was used in the optimized editing protocol (Figure 2B), and integration was scored by PCR and multiple positives were validated with sequencing across the junction as well as by microscopy. Strikingly, both types of hybrid dsDNA donor cocktails consistently yielded higher rates of accurate integration at three different loci, compared to melted and re-annealed traditional blunt donors (Figure 3C). We were successful at generating N- and C-terminal fusions with GFP and mCherry tags at rates comparable to ssODNs’, ~20% of F1 Rollers. Hybrid-B (Figure 3B) yielded the best precise editing rates, indicating that homology arms on the shorter product are not required to stimulate recombination.

## Discussion

CRISPR/Cas9 genome editing is highly efficient in *C. elegans* and should be accessible to investigators of all levels of experience. The protocols described here establish clear benchmarks for implementation and troubleshooting of *C. elegans* genome editing experiments. We demonstrate efficient editing at diverse genomic loci provided that editing targets are reasonably proximal (<20bp) to a PAM site. For short inserts (<140bp), we find the best efficiency with ssODN donors, as was previously reported (PAIX *et al.* 2015; PRIOR *et al.* 2017). For longer inserts, we recommend using donors that are hybrids of two asymmetric PCR products or a hybrid of a traditional symmetric donor and the insert (Figure 3B). The detailed version of our protocol is included in the supplemental materials (Appendix 2).

Our studies suggest several advantages of using a DNA-based gene expression marker for RNP-based editing. Perhaps most importantly, a plasmid-based expression marker reports on toxicity among a cohort of progeny that also inherit co-injected long dsDNA molecules. Long dsDNA molecules such as the pRF4 plasmid appear to traffic into a much smaller number of oocytes after injection into the syncytial ovary than do ssODN molecules. This likely explains why editing efficiencies with ssODN donors remain high at RNP concentrations that eliminate Roller progeny. The cohort of oocytes receiving sufficient plasmid DNA to express the Roller phenotype may also receive the highest amounts of RNP and thus are more sensitive to RNP-induced toxicity. Thus, the Roller phenotype identifies a portion of the brood that inherits longer dsDNA molecules, and thus is also likely to receive the co-injected long dsDNA donors required to serve as repair templates in some studies. In addition, the Roller phenotype is dominant and easily scored in the light dissecting microscope. Plasmid DNA preparation is inexpensive, and injection of plasmid DNA at these concentrations results primarily in transient F1 expression without further inheritance in subsequent generations. Indeed a recent study employed an *mCherry::myo-2* plasmid to identify ssODN templated editing events (PRIOR *et al.* 2017), demonstrating the feasibility of using other plasmid-based co-injection markers for genome editing. However, we find Roller more convenient as a fluorescence dissecting scope is not needed for scoring.

Our findings suggest that concentrations of Cas9 RNPs recommended in some protocols are quite toxic and eliminate F1 progeny that receive the largest amounts of co-injected long dsDNA. Since RNP activity and toxicity will vary depending on the specific target or guide sequence, or due to variations in protein preparation across vendors or home-made sources, we recommend that Cas9 RNPs be tested routinely for optimal concentration using the simple and inexpensive rol-6/TIDE approach (Figure 1A).

We do not yet know how or why hybrid dsDNA PCR donors stimulate HDR. It seems likely that other modifications, such as chemical modifications to the ends of the donor molecule may drive even greater efficiencies. However, we decided to report these findings now prior to fully exploring these issues, since the procedure for generating hybrid donors is extremely easy to implement and has worked efficiently on every one of over a dozen loci attempted in our group thus far. We anticipate that hybrid donors will also stimulate precise editing in other systems. In summary, it is now as easy to precisely edit the worm genome as it is to generate the iconic Roller transgenics first described by Mello et al. (MELLO *et al.* 1991). We strongly encourage even the total novice worm breeder to begin editing the genome of this fascinating “yeast” of metazoa.

## Acknowledgements

We thank Geraldine Seydoux at Johns Hopkins University and Scott Wolfe at the University of Massachusetts Medical School, Worcester for sharing Cas9 protein for pilot experiments. This work was funded by: American Cancer Society Fellowship (G.A.D.), Howard Hughes Medical Institute (C.C.M.), NIH PO1# HD078253 (C.C.M).

## Material and Methods

### Strains and Genetics

All the *C. elegans* strains were derived from Bristol N2 background and cultured on Normal Growth Media(NGM) plates seeded with OP50 bacteria (BRENNER 1974). Strains used in this study are listed in Table S1.

Sequences of all the oligos and crRNAs are provided in Appendix 1 and the detailed editing protocol is provided in Appendix 2.

Supplemental files available at FigShare and all the reagents are available upon request.

**Table S1:** List of *C. elegans* Strains

| Genotype | Strain Name |  |
| --- | --- | --- |
| glh-1(ne4595[gfp::glh-1])I | WM606 | In preparation |
| wago-4(ne4479[gfp::wago-4])II | WM610 | This study |
| sago-2(ne4480[gfp::sago-2])I | WM611 | This study |
| ppw-1(ne4481[gfp::ppw-1])I | WM609 | This study |
| gcna-1(ne4598[gfp::gcna-1])III | WM613 | This study |
| dvc-1(ne4599[mcherry::dvc-1])V | WM614 | This study |
| top-2(ne4600[top-2::mcherry])II | WM615 | This study |

#### Materials

1. S. *pyogenes* Cas9 3NLS (10 *μ*g/*μ*l, IDT)
2. tracrRNA (IDT*#* 1072532)
3. crRNA (2nmol or 10nmol, IDT)
4. ssDNA donor (standard desalting; 4 nmol Ultramer, IDT)
5. PRF4::*rol-6(su1006)* plasmid

#### Re-suspension

- Aliquot 0.5 *μ*l (5*μ*g) of Cas9 protein and store at −80%C to avoid freeze/thaw cycles. Use 1 aliquot per injection and add all the other reagents sequentially to the Cas9 tube.
- tracrRNA - 0.4 *μ*g/*μ*l in IDT duplex buffer, store at -20%C
- crRNA - 0.4 *μ*g/*μ*l in IDTE P.H 7.5, store at −20%C
- ssDNA oligo donor - 1 *μ*g/*μ*l in ddH2O, store at −20%C
- PRF4:: rol-6 *(su1006):* 500 ng/μl

## Donor Design and Generation

### ssODN donors

#### For generating short inserts (<130bp)

To generate a ssODN donor, add 35 bases of 5′ homology sequence in front of the tag and 35 bases of the 3′ homology sequence at the end. Remember to mutate the PAM site or the guide binding sequence if it is not already disrupted by the insert. If the guide binding sequence is mutated, length of homology sequence should be 35bp from the last mutation (PAIX *et al.* 2015).

#### For generating point mutations

Pick 35bp homology upstream and 35bp homology downstream of your guide cut site, which should ideally be within 20bp of the desired mutation site. Introduce the desired mutations in the donor and the PAM/ guide binding sequence.

#### For large deletions with 2 guides

Pick 35bp homology upstream of the left-guide cut site and 35bp homology downstream of the right-guide cut site and put them together. Everything in between will be removed. Deletions up to 1kb can be easily achieved through this strategy. In principle, this should work for larger deletions as well.

### dsDNA asymmetric-hybrid donors

1. Order 140bp oligos from IDT as Ultramers; 120bp as homology arms and 20bp complementary to GFP (or any other desired insert). Also, order standard oligos just complementary to your insert (no homology arms).
2. Generate two PCR products as shown below in figure; one with 120bp homology arms and the other without any homology arms (only insert sequence) using an insert containing plasmid as the template for PCR; perform 4 to 8 50 *μ*l reactions for each product.
3. Run 5 *μ*l on an agarose gel to check if a single bright band at ~1050bp (gfp +120bp+120bp) is obtained (in some cases the template plasmid band might be detected. It can be ignored as it does not interfere with HDR.). If non-specific amplification is observed, set up a temperature gradient and find the optimal temperature.
4. Combine all the PCR reactions of each product and column purify (we use Qiagen minElute kit), elute in 10-20 μl of water depending on brightness of the band, aiming to get >300ng/ul concentration.
5. Mix 1:1 of the purified PCRs (2μg:2μg for 20μl injection mix), heat to 95°C and cool to 4°C to re-anneal (95°C-2:00 min; 85°C-10 sec, 75°C-10 sec, 65°C-10 sec, 55°C-1:00 min, 45°-30 sec, 35°-10 sec, 25°-10 sec, 4°-forever.)
6. Add this donor cocktail to the rest of the injection mixture ONLY after pre-incubating Cas9, crRNA and tracrRNA (see below).

### Preparing injection mixtures

Add components of the injection mixture in the following sequence:

1. Cas9-0.5 μl of 10 μg/μl stock
2. Add tracrRNA - 5μl of 0.4 μg/μl stock
3. Add crRNA - 2.8μl of 0.4 μg/μl stock (if you are using two guides add 1.4 μl of each)
4. Incubate this mixture@37°C for 10 minutes before adding any DNA. Adding any double stranded DNA before RNP complex formation reduces HDR efficiency dramatically.
5. Add ssODN donor - 2.2 μl of 1μg/μl stock or dsDNA donor cocktail - 200 ng/μl (total 4 μg) in the final injection mixture
6. Add PRF4::rol-6 (*su1006*) plasmid - 1.6 μl of 500 ng/μl stock
7. If needed, add nuclease free water to bring the volume to 20μl
8. To avoid needle clogging, centrifuge the mixture@14000rpm for 2 min, transfer about 17 μl of the mixture to a fresh tube and proceed to loading the needles.

### Micro-injection and screening

1. Inject 10 to 20 animals and transfer them onto individual plates. After about 3 days, score for F1 rollers and place each roller onto a separate NGM plate.
2. In general, injections with ssODNs yield more number of F1 rollers per injected animal compared to the injections with dsDNA.

a. For ssODN-based editing: Pick 2 plates that segregate the most number of F1 rollers and from these 2 plates, pick about 24 F1 rollers and place them onto separate plates.
b. For dsDNA-based editing: Pick at least 24 F1 rollers from several plates and place them onto to separate plates.
3. To avoid false positives due to mosaicism in F1 animals, pick several F2s from each plate, perform lysis and genotyping. Genotyping primers should lie outside the homology arms to avoid false positives from transiently retained donor molecules. In some circumstances large inserts do not readily amplify in heterozygotes (because the small wild type band amplifies preferentially) In those situations it might be necessary to employ one primer inside the insert for each junction.
4. Alternatively, if the expression levels are detectable, insertions of fluorescent tags can be screened under a microscope either by using high magnification fluorescence microscope (mount several F2 animals onto 2% agarose pads) or by using a fluorescence dissecting scope.

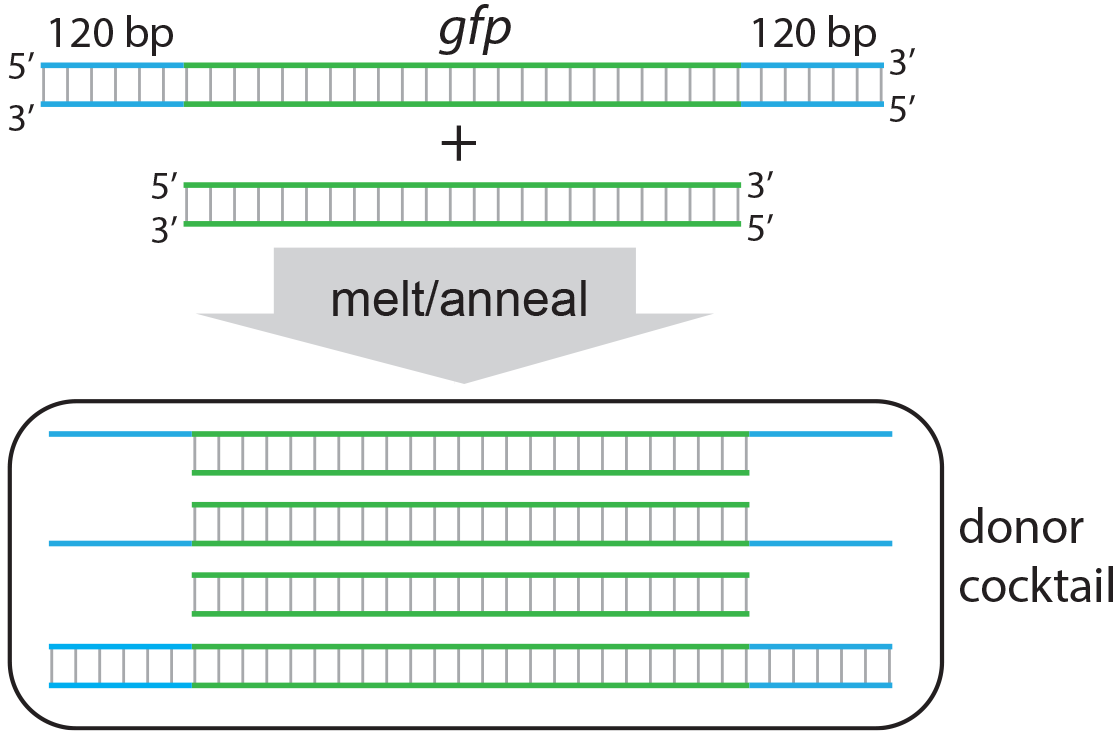

## Supplementary Figure Legends

**Figure S1.**
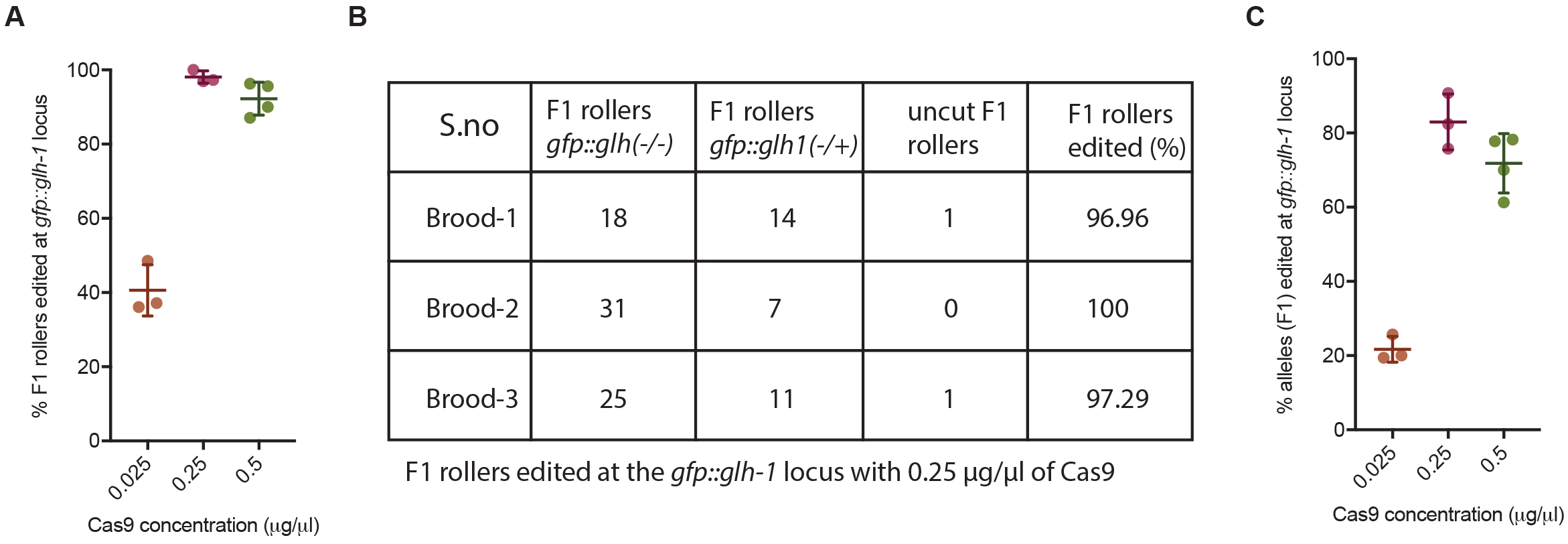
Monitoring editing efficiency at *gfp::glh-1* by microscopy. (**A**) Percentage of F1 rollers segregating GFP-negative F2 progeny plotted versus the concentration of Cas9 protein used in the injection mixture. (**B**) A break down of the F1 rollers among three of the broods from the 0.25μg/μl Cas9 injection that are homozygous or heterozygous for edits at *gfp::glh-1,* scored based on their F2 progeny. (**C**) Percentage of edited *gfp::glh-1* alleles calculated based on numbers of homozygous and heterozygous F1 rollers plotted versus the concentration of Cas9 protein used in the injection mixture (corresponds to TIDE data in **Figure 1C**). Error bars represent standard deviation from the mean.

**Figure S2.**
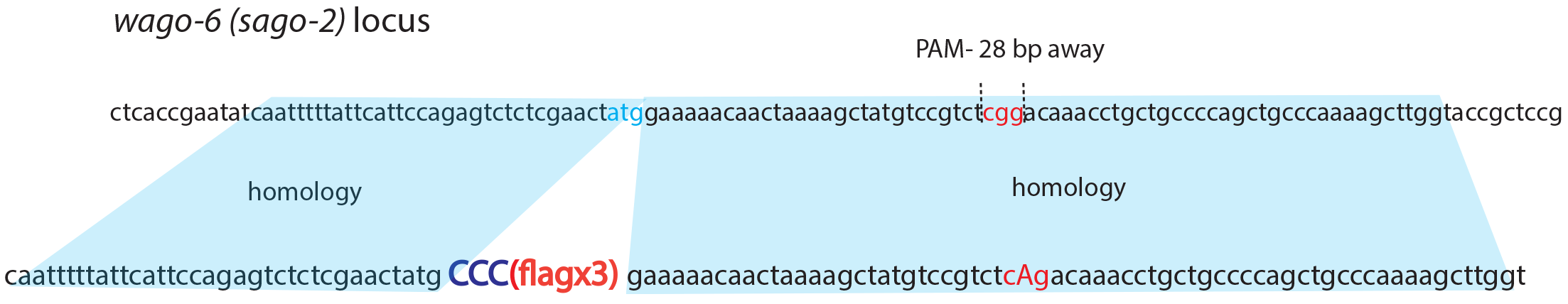
Alternative targeting strategy for *wago-6* (*sago-2*). A schematic representation of the donor design to introduce *flagx3* into the *wago-6* locus using the ATG-proximal PAM site that would not be normally disrupted by *flag* insertion. Briefly, introducing an extra proline codon (dark blue capital CCC) upstream of the flagx3 sequence in the donor for *wago-6* locus disrupts the PAM site in the donor.

